# TRIM24 preserves cardiomyocyte immune quiescence by repressing interferon/STAT signaling

**DOI:** 10.1101/2025.11.13.688187

**Authors:** Marco Neu, Ankush Borlepawar, Anushka Deshpande, Laura Neuberger, Tapan Kumar Baral, Frauke Senger, Elke Hammer, Nesrin Schmiedel, Michael Hausmann, Norbert Frey, Manju Kumari, Ashraf Yusuf Rangrez

## Abstract

Innate immune activation in cardiomyocytes is a key driver of inflammatory heart disease and heart failure progression, yet the mechanisms by which cardiomyocytes maintain transcriptional immune quiescence remain poorly understood. TRIM24, a multidomain chromatin reader implicated in cancer and inflammation, has not been previously studied in the heart. Here, we investigated whether TRIM24 regulates interferon/STAT-driven inflammatory programs and paracrine signaling in cardiomyocytes. Single-nucleus RNA sequencing revealed a pronounced downregulation of TRIM24 specifically in cardiomyocytes from human ischemic cardiomyopathy, a change not detected by bulk RNA-sequencing in either human or mouse myocardial infarction samples, likely due to masking by non-cardiomyocyte cell populations. Fractionated protein analysis further demonstrated that TRIM24 is enriched in cardiomyocyte nuclei. Functional studies in neonatal rat ventricular cardiomyocytes showed that TRIM24 represses interferon-stimulated and STAT-dependent genes, including Mx1, Irf7, and Ifit3. ChIP-Seq revealed TRIM24 occupancy at STAT-bound regulatory regions, suggesting cooperative suppression of inflammatory gene networks. Mechanistically, TRIM24 decreased Stat1a/b and Stat3 transcription, reduced protein abundance, and inhibited phosphorylation independently of proteasomal degradation; these effects were partially reversible by bromodomain inhibition. Functionally, TRIM24 overexpression dampened paracrine macrophage recruitment, whereas TRIM24 depletion amplified it. Collectively, these findings identify TRIM24 as a cardiomyocyte-enriched chromatin regulator that restrains STAT1/3 signaling and suppresses paracrine inflammatory activation. TRIM24 acts as a transcriptional safeguard of immune quiescence in the heart and represents a potential therapeutic target for limiting cardiac inflammation.

## Introduction

Cardiomyocytes rely on tightly regulated proteostasis to sustain contractile function and long-term survival ^1^. Protein homeostasis is maintained through molecular chaperones, autophagy, and the ubiquitin-proteasome system (UPS), which collectively ensure the clearance of damaged or misfolded proteins^1,2^. Beyond its canonical role in protein quality control, ubiquitination also serves as a versatile signaling modification that governs development, apoptosis and innate immune pathways, linking proteostasis to transcriptional networks that shape cardiomyocyte stress responses. Disruption of these networks can provoke proteotoxic stress and aberrant activation of inflammatory cascades, both of which are well-recognized drivers of cardiac pathology. Notably, the transcriptional circuitry that governs proteasome and UPS components in cardiomyocytes remains poorly defined; while transcription factors such as NFE2L1/NRF1 and NFE2L2/NRF2 have been shown to orchestrate proteasome and related stress-response genes in mammalian cells, and NRF1 has a documented role in cardiomyocyte proteostasis and regeneration^3,4^, the extent to which these and other chromatin regulators coordinate UPS gene networks and restrain inflammatory transcriptional programs in the heart is incompletely understood. Clarifying these transcriptional mechanisms is critical to understanding how cardiomyocytes maintain immune quiescence and to identifying targets to modulate maladaptive inflammation in heart disease.

The tripartite motif (TRIM) protein family, defined by a conserved RING-B-box-coiled coil (RBCC) architecture, includes numerous E3 ubiquitin ligases that integrates proteostatic regulation with cellular stress responses^5-7^. Several TRIM proteins have been implicated in cardiovascular biology, where they influence cardiac remodeling, hypertrophy, and inflammation. Among these, TRIM24, also known as transcription intermediary factor 1α (TIF1α), is of particular interest. TRIM24 is a multidomain protein that combines ubiquitin ligase activity with chromatin reader functions^8^. Through its C-terminal PHD finger and bromodomain, TRIM24 recognizes specific histone marks and recruits transcriptional cofactors, while its RING domain mediates ubiquitination of substrates such as p53^9-11^. Initially described as a nuclear receptor co-regulator in cancer, TRIM24 has since been implicated in both oncogenic and tumor-suppressive roles, depending on cellular context^12,13^.

TRIM24 has emerged as a versatile regulator of immune response across diverse disease models. TRIM24-deficient mice exhibit heightened STAT1^14^, while elevated TRIM24 expression has been reported in inflammatory disorders such as dermatomyositis^15^. Similarly, TRIM24 loss enhances IL-10 expression via the JAK/STAT3 axis, conferring resistance to endotoxic shock^16^. Mechanistically, TRIM24 suppresses interferon pathways by cooperating with retinoic acid receptor α (RARα) at the Stat1 promoter, thereby limiting downstream activation of IRF7 and guanylate-binding proteins^17^. Yet TRIM24’s immune-regulatory functions are context-dependent. In macrophages and tumor models, its depletion impairs interferon-stimulated gene expression and enhances viral replication^18^. TRIM24 also interacts with JAK2 and STAT3, supports Th2 cell activation, and regulates chemokine expression^19,20^.Moreover, it modulates STAT6 acetylation in macrophages, influencing polarization toward pro- or anti-inflammatory states^21^. Collectively, these findings highlight TRIM24 as a versatile regulator of immune signaling and cellular homeostasis.

In the heart, TRIM24 expression is elevated in the diseased myocardium, where it promotes hypertrophy by stabilizing dysbindin and counteracting TRIM32-mediated degradation^22^. More recently, our group demonstrated that TRIM24 regulates chromatin remodeling and calcium dynamics in cardiomyocytes, thereby directly influencing cardiac excitation-contraction coupling^23^. However, despite its growing recognition as a chromatin-associated modulator of stress responses, its role in coordinating immune transcriptional networks within cardiomyocytes remains unexplored. This gap is particularly striking given that innate immune activation in cardiomyocytes is a key driver of inflammatory cardiomyopathies and heart failure progression, yet the transcriptional safeguards that preserve immune quiescence under physiological conditions remain poorly defined. Emerging single-nucleus transcriptomic data indicate that TRIM24 is among the most enriched chromatin regulators in healthy cardiomyocyte nuclei but is markedly reduced in failing human hearts and after myocardial infarction. These observations position TRIM24 as a potential transcriptional gatekeeper linking chromatin state to immune homeostasis. Given that cardiomyocytes are not passive bystanders but active participants in innate immune signaling, defining how TRIM24 modulates their inflammatory responses is crucial to understanding cardiac resilience to stress.

We, therefore, hypothesized that TRIM24 functions as a chromatin-mediated brake on interferon-driven signaling in cardiomyocytes, thereby preserving immune quiescence and proteostatic stability. In the present study, we employed integrative transcriptomic, proteomic, and chromatin profiling combined with functional assays in neonatal rat ventricular cardiomyocytes to dissect the immunomodulatory role of TRIM24. We show that TRIM24 represses interferon-stimulated genes and STAT1/3 signaling, reduces macrophage recruitment through paracrine mechanisms, and exerts its effects via its chromatin reader domain rather than its E3 ligase activity. By integrating human single-nucleus datasets, in vivo myocardial injury models, and mechanistic in vitro studies, we identify TRIM24 as a nuclear safeguard that restrains inflammatory signaling in cardiomyocytes and preserves cardiac immune homeostasis.

## Materials and Methods

### Animal experiments

All animal procedures were approved by the Animal Welfare Officer at the Ministry of Agriculture, Rural Areas, Europe and Consumer Protection (MLLEV), Schleswig-Holstein, and conducted in compliance with institutional ethical standards. Mice were maintained at 23 °C under a 12 hours light/dark cycle with free access to standard chow (Altromin 1329P) and water.

### Left anterior descending (LAD) coronary artery ligation–induced myocardial infarction

Acute (3-day) and chronic (30-day) myocardial infarction (MI) were induced by permanent ligation of the left anterior descending (LAD) coronary artery. Mice received pre-operative analgesia with buprenorphine (0.1 mg/kg, s.c., Temgesic®). General anesthesia was induced and maintained with isoflurane (3% for induction, delivered in oxygen). Animals were intubated with a flexible catheter and mechanically ventilated (tidal volume 200 μL; 120–130 breaths/min). Local anesthesia at the thoracotomy site was performed by subcutaneous and intramuscular injection of 2% lidocaine (0.2–0.3 mL). A left thoracotomy was performed at the third intercostal space. After retraction of the fourth and fifth ribs, the pericardium was opened and the heart was gently exteriorized. The LAD was identified and permanently ligated using an 8-0 non-absorbable monofilament suture (Prolene®). The heart was then returned to the thoracic cavity, the chest wall was closed, and negative pressure was restored by evacuation of intrathoracic air. The skin was closed with 4-0 Vicryl Rapid® (Ethicon®). During surgery and recovery, body temperature was continuously monitored using a rectal probe and maintained via a heated platform with automatic feedback control. Mice were placed in individual cages for undisturbed recovery and received post-operative analgesia for 2–3 days via tramadol-supplemented drinking water (2.5 µg/mL, ad libitum). Animals were euthanized and tissues collected either 3 days post-LAD ligation (acute MI) or 30 days post-LAD ligation (chronic MI) for downstream analyses.

### Cloning of TRIM24 overexpression and knockdown constructs

Adenoviral constructs for TRIM24 overexpression and knockdown were generated based on the protocol described by Borlepawar et al. (2017)^22^. The expression plasmid containing TRIM24 was kindly provided by Dr. David Barton (University of Texas, Houston). For TRIM24 knockdown, synthetic microRNAs were designed and cloned using the BLOCK-iT™ Pol II miR RNAi Expression System in combination with the Gateway® cloning platform (Thermo Fisher Scientific, Waltham, MA, USA). To generate adenoviruses expressing full-length human TRIM24, the ViraPower™ Adenoviral Expression System (Thermo Fisher Scientific) was used following the manufacturer’s instructions. Specifically, TRIM24 cDNA previously cloned into the pDONR221 entry vector was recombined into the pAd/CMV/V5-DEST destination vector. The resulting plasmids were linearized using PacI and subsequently transfected into HEK293A cells to produce viral particles. Viral titers were determined by infecting HEK293A cells and detecting expression with a fluorescent anti-Hexon antibody. An adenoviral construct encoding β-galactosidase-V5 (Ad-LacZ; Thermo Fisher Scientific) was used as a control.

### Neonatal rat ventricular cardiomyocyte (NRVCM) isolation and culture

Primary neonatal rat ventricular cardiomyocytes (NRVCMs) were isolated from the left ventricles of 1-2-day-old Wistar rat pups (Charles River, Lyon, France) following a protocol modified from Borlepawar et al. (2017) ^22^. The excised ventricles were finely chopped and incubated in ADS buffer (120 mmol/L NaCl, 20 mmol/L HEPES, 8 mmol/L NaH_2_PO_4_, 6 mmol/L glucose, 5 mmol/L KCl, 0.8 mmol/L MgSO_4_, pH 7.4). Enzymatic digestion was carried out in five to six cycles using pancreatin (0.6 mg/mL; Sigma) at 37°C, followed by treatment with collagenase type II (0.5 mg/mL; Worthington, Lakewood, NJ, USA) in sterile ADS buffer. To stop digestion, the cell suspension was filtered and supplemented with newborn calf serum. Cardiac fibroblasts were removed by density gradient centrifugation using Percoll (GE Healthcare, Chicago, IL, USA). The resulting cardiomyocyte fraction was cultured in DMEM supplemented with 10% fetal calf serum, 2 mM L-glutamine, and penicillin/streptomycin (PAA Laboratories, Pasching, Austria). Adenoviral transduction was performed 24 hours after plating: cells were infected with LacZ or TRIM24 adenovirus (MOI 25) for overexpression, or with mir-Neg or mir-TRIM24 (MOI 500) for knockdown experiments. Infections were carried out in serum-free DMEM containing antibiotics, and cells were harvested 72 hours later for further analyses.

### Isolation and cell fractionation of adult cardiomyocytes

Mouse adult ventricular cardiomyocytes were isolated using a Langendorff-independent isolation procedure as described by Foo and colleagues^24^. After isolation, cells were kept on ice and processed immediately to minimize protein degradation. Subcellular fractionation into cytoplasmic, membrane-associated, and nuclear fractions was performed following the methodology reported by Baghirova et al.^25^. Briefly, cardiomyocytes were resuspended in a mild lysis buffer formulated to disrupt the plasma membrane without solubilizing intracellular organelles. Mechanical disruption was applied to aid cell lysis. After lysis, samples underwent a series of differential centrifugation steps to sequentially separate the cytoplasmic fraction from heavier membrane-containing components. The supernatant from this initial centrifugation constituted the cytoplasmic fraction. The remaining pellet was resuspended in a membrane extraction buffer containing detergents optimized to selectively release membrane-associated and organelle-bound proteins. Insoluble nuclear material was separated from this extract by further centrifugation. The final pellet, consisting primarily of nuclei, was solubilized in a high-stringency buffer to obtain the nuclear protein fraction. Protein concentration of each fraction was quantified using the bicinchoninic acid (BCA) assay. Purity of the fractions was assessed by immunoblotting using well-established marker proteins: GAPDH for the cytoplasmic fraction, VDAC (voltage-dependent anion channel protein) for the membrane-associated fraction, and Histone H3 for the nuclear fraction.

### Immunostaining

NRVCMs were fixed in 4% paraformaldehyde (PFA) for 10 minutes at 37°C, followed by several washes with phosphate-buffered saline (PBS). To block unspecific binding sites, cells were incubated for 1 hour at room temperature in 2.5% bovine serum albumin (BSA) prepared in PBS. Primary antibody staining was performed by incubating the cells overnight at 4°C in the same blocking solution containing the appropriate primary antibodies at their respective dilutions. On the following day, cells were washed thoroughly with PBS (five to six times) and subsequently incubated overnight at 4°C with the corresponding secondary antibodies, also diluted in blocking buffer. After another series of PBS washes, nuclei were counterstained with DAPI (D1306, Invitrogen, Waltham, MA, USA) at a 1:10,000 dilution for 1 hour at room temperature. Confocal imaging was carried out using the LSM980 AIRY system (Zeiss, Oberkochen, Germany).

### Localization microscopy

The custom-built microscope used for super-resolution imaging was located at the Light Microscopy Facility of the German Cancer Research Center (DKFZ) in Heidelberg and constructed by W. Schaufler. A detailed technical description of the setup is available elsewhere^26-28^. To maintain consistent imaging conditions, the system was designed with enhanced thermo-mechanical stability, enabling a localization precision of ±10 nm. Alexa Fluor 647 was excited using a 642 nm laser at an intensity of 0.8 kW/cm^2^ at the sample plane, while Alexa Fluor 594 excitation was achieved using a 561 nm laser at 1.87 kW/cm^2^. Each cell was imaged for 2000 frames with an exposure time of 100 ms per frame. Super-resolution signal coordinates were determined using a proprietary software that detects intensity variations between consecutive frames. Fluorophores entering dark states for more than two frames were identified, and their signal barycenters were computed. Only signals above a defined intensity threshold were retained and recorded in the “Orte-Matrix”^26^.

For spatial distribution analysis, Ripley’s K-function was employed to compute and normalize point-to-point distances^29,30^. To characterize the topological features of the signal patterns, persistent homology analysis was conducted as described previously^29,30^. This method identifies and quantifies structural features in the data by transforming spatial distributions into a topological space. Two key parameters were extracted: the number of components (representing the initial points) and the number of holes (corresponding to enclosed spaces formed during the analysis). Each point in the Orte-Matrix was treated as the center of an expanding virtual circle. Bars in the persistence diagram originated at radius zero (one component) and continued to grow until two circles overlapped, at which point one bar terminated as the components merged. As more circles intersected, closed loops, or “holes”, formed. These were also represented as bars, beginning with hole formation and ending once the hole was entirely covered by overlapping circles. The frequency of bar endpoint values was used as a proxy for hole size distribution.

### Antibodies

#### Antibodies used for various experiments in this study were as follows

Flag-tag, rabbit/mono, Merck, F2555, ChIP (1:1000); GAPDH, rabbit/mono, Cell Signaling, 2118S, WB (1:1000); TRIM24, rabbit/poly, Proteintech, 14208-1-AP, IF/WB (1:1000); Vinculin, rabbit/mono, Cell Signaling, 4650S, WB (1:1000) Flag-tag, rabbit, mono, Merck, ChIP (1:10,000); STAT1, rabbit/mono, Cell Signaling, 14994S, IF/WB (1:1000); phospho-STAT1, rabbit/mono, Cell Signaling, 9167S, IF (1:500)/WB (1:1000); STAT3, rabbit/mono, Cell Signaling, 12640S, IF/WB (1:1000); phosphor-STAT3, rabbit/mono, Cell Signaling, 9145S, IF (1:500)/ WB (1:1000); AlexaFLuor 546, anti-rabbit/anti-mouse, donkey, Invitrogen, A10041/A10036, (1:2000).

### RNA-sequencing

Library preparation, RNA sequencing, and bioinformatic analyses were outsourced to Arraystar Inc. Go Beyond RNA (Rockville, MD, USA) and performed as previously described^31^. Total RNA from each sample was quantified using a NanoDrop ND-1000 spectrophotometer. For library construction, approximately 2 µg of total RNA per sample was processed. First, polyadenylated mRNA was isolated using oligo(dT) magnetic beads to deplete ribosomal RNA. Subsequently, strand-specific RNA-seq libraries were prepared using the KAPA Stranded RNA-Seq Library Preparation Kit (Illumina, San Diego, CA, USA), which incorporates dUTP during second-strand synthesis to preserve strand orientation. Library quality was assessed using the Agilent 2100 Bioanalyzer (Agilent, Waldbronn, Germany), and concentrations were determined via absolute quantification qPCR. Indexed libraries were pooled and denatured with NaOH to generate single-stranded DNA, which was captured on an Illumina HiSeq 4000 flow cell, amplified on-chip, and sequenced in paired-end mode with 150 cycles per read.

Raw sequencing data were processed using the Solexa pipeline (v1.8) and quality-checked using FastQC (http://www.bioinformatics.babraham.ac.uk/projects/fastqc/). Adapter sequences were removed using Cutadapt, and reads were aligned to the reference genome using Hisat2. Transcript assembly and quantification were performed with StringTie, and expression levels were calculated as FPKM values using the Ballgown package in R. Differential gene and transcript expression analyses were conducted with Ballgown. Novel transcripts were identified by comparing assembled transcripts to the reference annotation, and their coding potential was evaluated with CPAT. Downstream statistical and exploratory analyses, including principal component analysis (PCA), correlation matrices, hierarchical clustering, gene ontology (GO) analysis, KEGG pathway enrichment, scatter plots, and volcano plots, were conducted using R, Python, or shell-based workflows. GO and KEGG enrichment analyses were performed via shinyGO 0.82 (South Dakota State University, USA) using an FDR threshold of 0.05. All RNA-seq-related source data, including those corresponding to the figures in the main manuscript, are available in Supplementary Data Table 1.

### Single-cell Data Integration and Clustering

All analyses were performed using R version 4.5.1 (2025-06-13) and Seurat v5.3.0. For GSE161921, cells with less than 200 or more than 2000 detected genes were filtered out, as well as genes expressed in fewer than 3 cells. Mitochondrial genes were removed, and each sample was normalized with SCTransform, then integrated together using Seurat’s SCT-based integration workflow. Dimensionality reduction was performed via PCA, and clustering done at a range of resolutions. Final clusters were annotated using module scores that were computed from curated marker genes for major cardiac cell types, in using methods described in^32^.

For GSE121893, similar filtering thresholds were applied (min. 200 genes per cell, min. 3 cells per gene) and normalization was performed using SCTransform. To analyze cardiomyocyte subtypes, we merged only the cells annotated as Cardiomyocytes from GSE161921 with all cells from GSE121893. Batch correction was performed using Harmony, and downstream analysis (PCA, UMAP, clustering) was performed in the Harmony-corrected space.

### Proteomics analysis

Four biological replicates from independent experiments were analyzed per experimental condition. Protein extraction was performed using a urea/thiourea buffer (8 M/2 M), combined with repeated freeze-thaw cycles. Following nucleic acid fragmentation via sonication, lysates were centrifuged for 60 minutes at 20 °C and 16,000 × g to collect the protein-containing supernatant. For each sample, 3 µg of protein was subjected to reduction with dithiothreitol and alkylation with iodoacetamide, followed by enzymatic digestion using trypsin at a protein-to-enzyme ratio of 25:1. Digestion proceeded for 16 hours at 37 °C. The resulting peptide mixtures were desalted using C18 µZipTip pipette tips (Merck Millipore, Darmstadt, Germany).

Mass spectrometry was conducted on a nano-Acquity UPLC Synapt G2-Si HDMS system (Waters, Milford, MA, USA), as previously described ^33^. Peptide and protein identification was performed using PLGS software version 3.3 (Waters), with reference to the NCBI RefSeq FASTA database (*Rattus norvegicus* version 03_2019; 66,850 sequences). Search parameters included trypsin as the proteolytic enzyme (allowing one missed cleavage), carbamidomethylation of cysteine as a fixed modification, and oxidation of methionine as a variable modification. Identified proteins were further analyzed using ISOQuant version 1.8 ^34^. Quantification was based on the three most intense, unmodified peptides per protein (minimum signal intensity: 3,000). For normalization, systematic errors were corrected using the LOWESS method with a bandwidth setting of 0.3 ^35^. Group comparisons were performed using Student’s t-test, and differences were considered statistically significant at *p* < 0.05.

### Protein preparation and immunoblotting

Seventy-two hours following adenoviral transduction, NRVCMs were lysed using three freeze-thaw cycles in RIPA buffer (20 mM Tris, 10 mM DTT, 500 mM NaCl, 1% NP-40, 12.5% glycerol), supplemented with protease and phosphatase inhibitor cocktails (Roche, Mannheim, Germany). Cell debris was removed by centrifugation at 18,000 × *g* for 20 minutes. Protein concentrations in the cleared lysates were measured using the DC Protein Assay Kit (Bio-Rad Laboratories, San Francisco, CA, USA). Equal amounts of total protein were separated via SDS-PAGE on either standard 10% polyacrylamide gels or pre-cast 4-12% gradient gels (Life Technologies, Carlsbad, CA, USA). Proteins were transferred to nitrocellulose membranes and blocked for 1 hour at room temperature with 5% non-fat dry milk dissolved in 0.1% TBST (Tris-buffered saline containing Tween-20). Membranes were incubated overnight at 4 °C with primary antibodies diluted in blocking buffer. The following day, membranes were washed four times in TBST and subsequently incubated with HRP-conjugated secondary antibodies (1:10,000; Cell Signaling, Danvers, MA, USA). Signal detection was performed using the ECL Select chemiluminescence kit (Merck, Darmstadt, Germany), and chemiluminescent signals were visualized using a ChemiDoc imaging system (Bio-Rad, San Francisco, CA, USA). Densitometric analysis of protein bands was conducted using ImageJ/Fiji software (version 1.46).

### RNA isolation and qPCR

Total RNA was isolated from NRVCMs using TRIzol reagent (Invitrogen, Waltham, MA, USA) following the manufacturer’s instructions. To generate complementary DNA (cDNA), 1 µg of DNase-treated total RNA was reverse-transcribed using the LunaScript First-Strand cDNA Synthesis Kit (New England Biolabs GmbH, Frankfurt, Germany). Quantitative real-time PCR (qRT-PCR) was conducted using the EXPRESS SYBR Green ER reagent (Life Technologies, Carlsbad, CA, USA) on a LightCycler 480 II system (Roche, Mannheim, Germany). The PCR protocol consisted of an initial denaturation step at 95 °C for 3 minutes, followed by 40 amplification cycles (15 seconds at 95 °C, 45 seconds at 60 °C). Gene expression levels were normalized to Rpl32 as a housekeeping gene. Primer sequences are listed in Supplementary Table 1. All NRVCM-based experiments were conducted in six technical replicates and independently repeated three times.

### Macrophage migration Assay

Conditioned medium from interferon-stimulated NRVCMs was harvested and applied to the lower compartment of an 8 μm pore Transwell system. In the upper chamber, RAW 264.7 macrophages (1×10^5^ cells/well) were seeded and incubated at 37°C for 24 hours to assess chemotactic migration. After incubation, non-migrated cells were removed from the upper membrane surface using a cotton swab. Cells that had migrated to the lower membrane side were fixed in 100% methanol for 10 minutes and stained with 0.1% crystal blue in 0.1 M borate buffer for 20 minutes at room temperature. Migrated cells were imaged using an inverted microscope. For quantification, membranes were excised and incubated in 10% acetic acid (200 μl) to extract the dye, and absorbance was measured at 560 nm.

### Chemical treatments

To modulate signaling pathways and mimic pathological or physiological stress conditions, neonatal rat ventricular cardiomyocytes (NRVCMs) were treated with specific chemical agents and inhibitors at defined concentrations and incubation times. MG132 (10 μM, 12 hours) was used to inhibit proteasomal degradation and thereby interfere with UPS-mediated protein turnover. Recombinant interferon γ (IFNγ, 75 ng/ml, 24 hours) was applied to stimulate type I interferon signaling. Lipopolysaccharide (LPS, 100 ng/ml, 12 hours) served to activate innate immune pathways via toll-like receptor 4. The selective bromodomain inhibitor IACS-9571 was used at concentrations ranging of 50 nM for 24 hours to modulate epigenetic regulations of TRIM24. All treatments were performed under standard cell culture conditions, and controls received vehicle-only solutions.

### Chromatin immunoprecipitation

Neonatal cardiomyocytes were washed twice with PBS and fixed with 1% (v/v) formaldehyde for 10 minutes on ice to crosslink proteins and DNA. Crosslinking was quenched by the addition of glycine to a final concentration of 250 mM for 5 minutes. Cells were then snap-frozen in liquid nitrogen and stored until further processing. Chromatin extraction was carried out using a sequential lysis protocol with three different buffers. Cells were incubated for 10 minutes at 4 °C under constant rotation with Lysis Buffer 1 (50 mM Hepes-KOH pH 7.5, 140 mM NaCl, 1 mM EDTA, 10% glycerol, 0.5% NP-40, 0.25% Triton X-100), followed by Lysis Buffer 2 (200 mM NaCl, 1 mM EDTA pH 8, 0.5 mM EGTA pH 8, 10 mM Tris pH 8) for another 10 minutes. Cells were then transferred to Lysis Buffer 3 (50 mM Tris pH 8, 0.1% SDS, 0.95% NP-40, 0.1% sodium deoxycholate, 10 mM EDTA, 150 mM NaCl) and sonicated in polymethylpentene (TPX) tubes (Diagenode, Seraing, Belgium) using the Bioruptor automated sonication system (Diagenode). Sonication was performed at high intensity for 40 cycles (30 seconds on, 30 seconds off).

Chromatin immunoprecipitation (ChIP) was conducted using the semi-automated IP-Star system (Diagenode), following the manufacturer’s instructions. All buffers and magnetic beads were purchased from Diagenode. For each pulldown, 1 µg of chromatin was incubated with anti-FLAG antibody. Chromatin samples were then treated with RNase (Qiagen, Venlo, Netherlands) and Proteinase K (Roche, Mannheim, Germany), followed by overnight reverse crosslinking at 65 °C. DNA concentrations were measured using the Qubit® dsDNA High Sensitivity Assay Kit (Thermo Fisher Scientific). DNA was subsequently ethanol-precipitated and quality was assessed by agarose gel electrophoresis. Library preparation was performed using the NEBNext DNA Library Prep Master Mix Set for Illumina (E6040) according to the manufacturer’s protocol. Sequencing was carried out by Novogene (Cambridge, UK). Downstream analysis was performed using shinyGO 0.82 (South Dakota State University, USA), applying a false discovery rate (FDR) threshold of 0.05. ChIP-seq data are available under accession number E-MTAB-8011.

### Statistical Analysis

All results are shown as the means ± SEM. The statistical analyses of the data were performed using a two-tailed Student’s *t-*test. p-values of less than 0.05 were considered statistically significant.

## Results

### TRIM24 is a cardiomyocyte-enriched nuclear factor reduced in human ischemic cardiomyopathy and regionally regulated after myocardial infarction

Innate immune activation within cardiomyocytes is increasingly recognized as a major driver of inflammatory heart disease, yet the transcriptional mechanisms that maintain immune homeostasis in the myocardium are poorly defined. TRIM24 is a multidomain transcriptional co-regulator and E3 ubiquitin ligase that integrates chromatin state with transcription factor activity through its PHD–bromodomain module. Although TRIM24 modulates immune and stress signaling in non-cardiac tissues, its function in cardiomyocytes has not been examined. To examine TRIM24 expression in the context of inflammation-driven cardiac pathology, we analyzed a high-resolution snRNA-seq dataset comprising >99,000 nuclei from non-infarcted left ventricular tissue of ischemic cardiomyopathy (ICM) transplant recipients and non-failing donor controls generated by the Ellinor group^36^. TRIM24 transcripts were present in several cardiac cell types, but were most abundant in cardiomyocytes (Figure 1A, 1B). A dot-plot summary indicates that though the Trim24 transcript levels are not significantly altered in cardiomyocytes, it is mildly upregulated in cardiac lymphatic endothelial cells, mast cells, and proliferating macrophages in ICM compared to non-failing (NF) hearts (Figure 1C). Consistent with these cell-resolved results, bulk RNA-seq from whole human hearts (GSE141910) did not detect significant differences in TRIM24 expression between ICM and non-failing controls (Supplementary Figure 1A), Interestingly however, analysis of an independent patient snRNA-seq dataset^37^ revealed reduced TRIM24 expression in cardiomyocytes and endothelial cells in heart failure, with mild increases in fibroblasts, macrophages, and smooth muscle cells (Supplementary Figure 1B, 1C).

**Figure 1.**
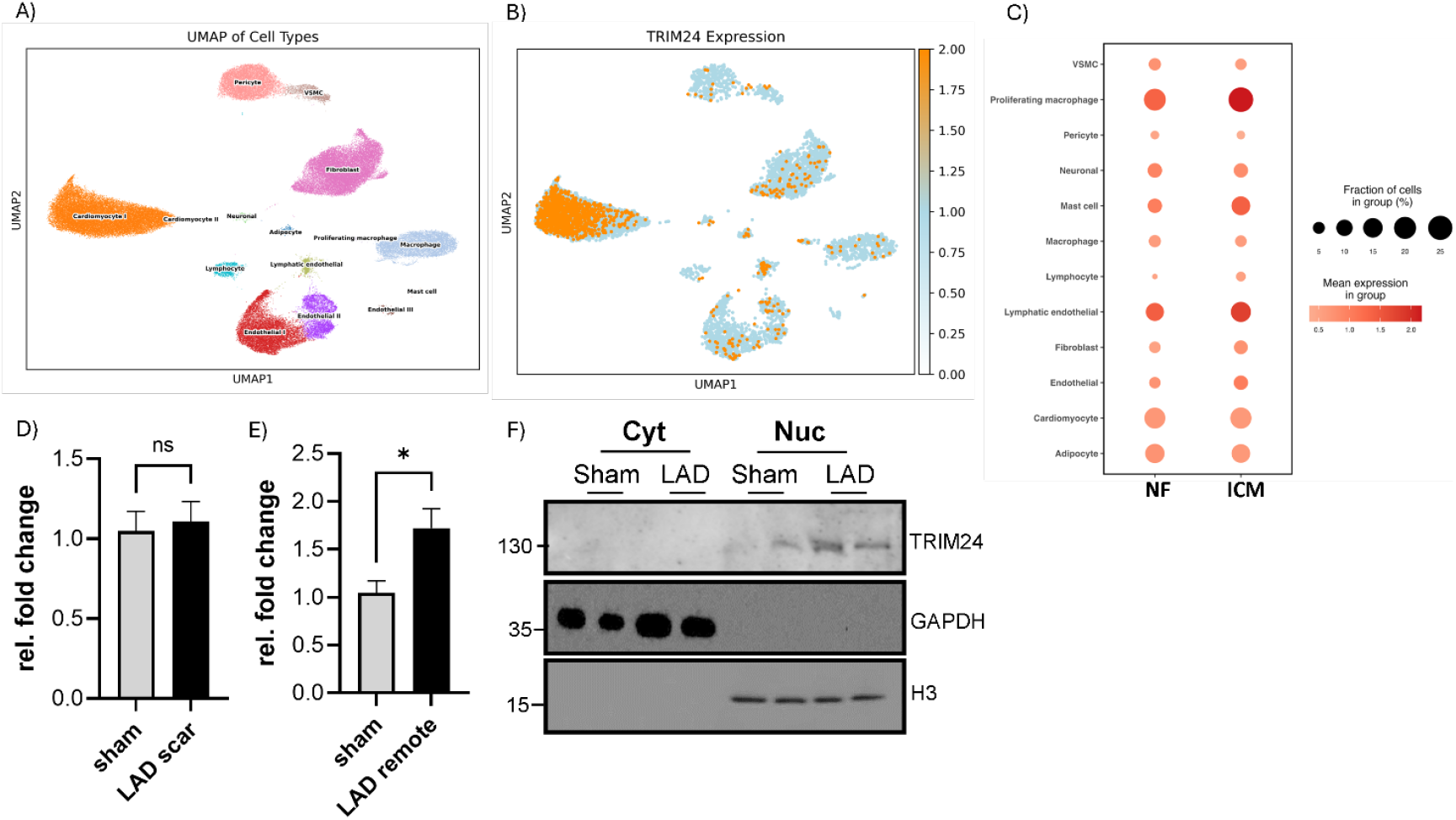
TRIM24 is a cardiomyocyte-enriched nuclear factor reduced in human ischemic cardiomyopathy and regionally regulated after myocardial infarction. UMAP plot showing cell types identified (**A**) and TRIM24 expression (highlighted in orange) across major cardiac cell populations in human left ventricle single-nucleus RNA sequencing dataset (Broad Institute’s Single Cell Portal SCP1849), demonstrating highest expression in cardiomyocytes. Quantification of Trim24 transcripts from mouse hearts after left anterior descending coronary artery (LAD) ligation compared with sham controls, showing unchanged levels in the scar region (**D**) and upregulation in the remote region (**E**). (**F**) Subcellular localization of TRIM24 protein in cardiomyocytes isolated from sham and LAD-operated mouse hearts, showing predominant nuclear enrichment with minimal cytoplasmic signal. Statistical significance was determined using two-tailed Student’s t test. Error bars represent mean ± SEM. ns, non-significant; *, *p* < 0.05.

Additionally, to assess regulation of TRIM24 under cardiac injury, we analyzed TRIM24 expression level in the mouse left anterior descending (LAD) coronary ligation model. Trim24 transcript abundance was unchanged in the infarct (scar) region but significantly upregulated in the remote myocardium compared with sham controls (Figure 1D, 1E). In contrast, bulk RNA-seq data from whole human (GSE141910) and mouse hearts showed no significant differences in TRIM24 expression in ICM (Supplementary Figure 1C-E), likely reflecting dilution of cardiomyocyte-specific signals in whole-tissue homogenates. Interestingly, acute systemic inflammatory stress led to the opposite effect: in LPS-treated mouse hearts (GSE185754), Trim24 transcript levels were significantly reduced within 24 h (Supplementary Figure 1F). To further define the cellular and molecular context in which TRIM24 operates in the heart, we examined its subcellular localization in adult mouse cardiomyocytes isolated from sham and LAD-induced ischemic mice. Subcellular fractionation revealed that TRIM24 is predominantly nuclear under both conditions, with moderate enrichment in the nuclear fraction following LAD ligation and negligible presence in the cytoplasm (Figure 1F). Extended cell fractionation including total lysate and membrane fractions confirmed that TRIM24 is abundant in nuclear fractions, weakly detected in membrane fractions, and virtually absent from the cytoplasm (Supplementary Figure 1G). This nuclear confinement is consistent with its predicted role as a chromatin-associated regulator of gene expression.

Taken together, these analyses identify TRIM24 as a cardiomyocyte-enriched nuclear transcriptional co-activator that is downregulated in human ICM, regionally upregulated in the remote myocardium after infarction, and suppressed during systemic inflammation. These findings provide physiological and pathological context dependent role for TRIM24’s in maintaining cardiomyocyte homeostasis and restraining inflammatory transcriptional programs described in subsequent sections.

### TRIM24 reprograms the cardiomyocyte immune transcriptome and targets STAT-related regulatory networks

Having established that TRIM24 is a cardiomyocyte-enriched nuclear factor dynamically regulated in cardiac injury, we next investigated its functional impact on cardiomyocyte gene expression. To this end, we performed RNA sequencing in neonatal rat ventricular cardiomyocytes (NRVCMs) overexpressing TRIM24. This analysis identified 266 differentially expressed genes, of which 206 were downregulated and 60 upregulated, indicating a predominantly repressive role for TRIM24. Enrichment analyses (GSEA, GO, KEGG) revealed strong suppression of immune-related pathways, including type I and II interferon signaling, cytokine-mediated signaling, and antiviral defense modules such as viral myocarditis and RIG-I-like receptor signaling (Figure 2A, 2B; Supplementary Figure 2A-L). Among the most consistently repressed genes were *Mx1, Irf7*, and *Ifit3*, hallmark interferon-stimulated genes involved in viral recognition, inflammasome activation, and amplification of cytokine signaling through the STAT axis (Figure 2A-C). Furthermore, proteomics analysis of NRVCMs after TRIM24 knockdown revealed increased levels of proteins involved in STAT signaling, including STAT1 (Supplementary Figure 2M).

**Figure 2.**
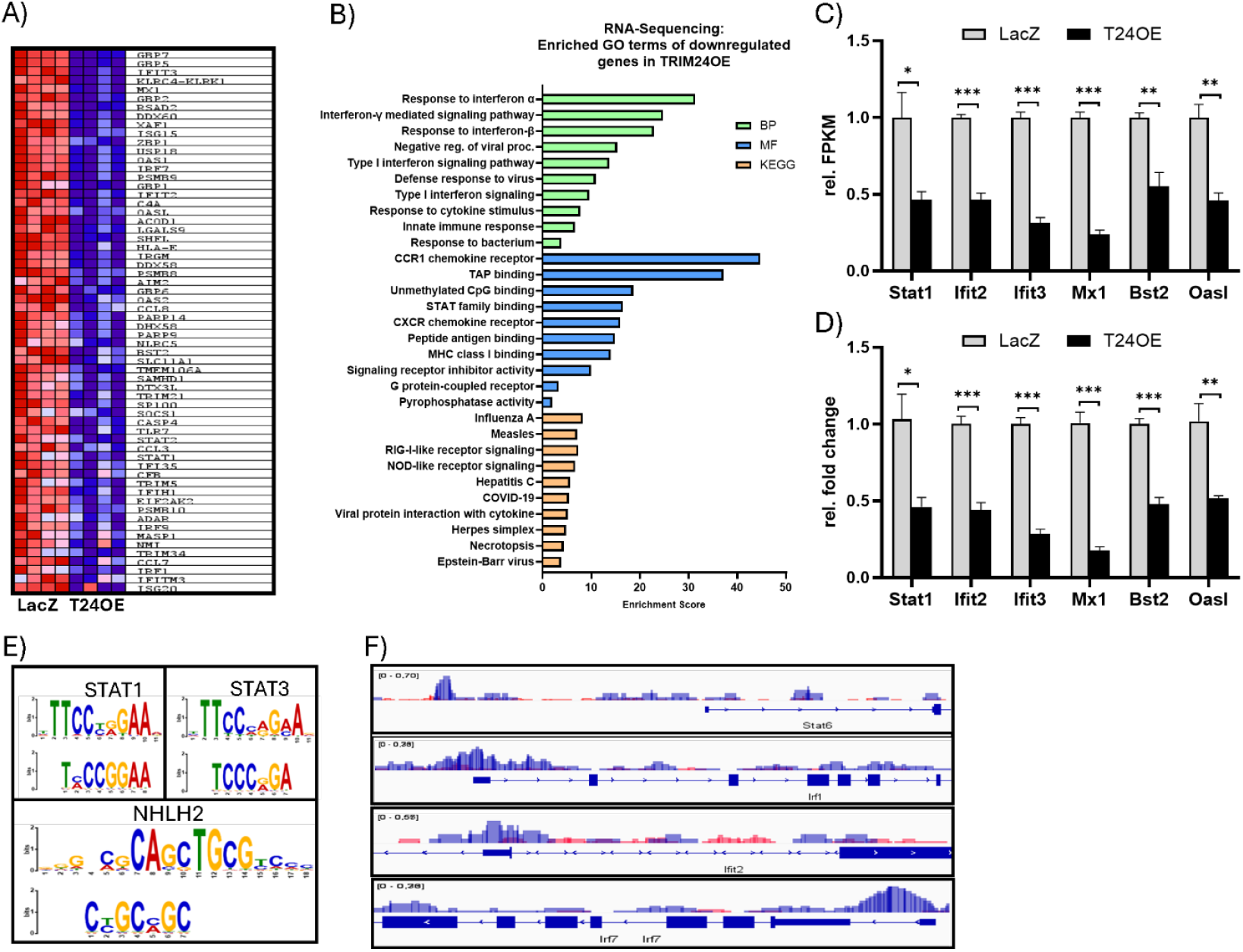
TRIM24 reprograms the cardiomyocyte immune transcriptome and targets STAT-related regulatory networks. (**A**) Heatmap of differentially expressed genes in neonatal rat ventricular cardiomyocytes (NRVCMs) overexpressing TRIM24 (T24OE) compared with LacZ controls, highlighting strong downregulation of interferon-stimulated genes, including, *Mx1, Irf7*, and *Ifit3*. (**B**) Functional enrichment analysis of downregulated transcripts showing suppression of immune-related Gene Ontology (GO) categories, including response to type I and type II interferon signaling, cytokine-mediated signaling, STAT-family binding. (**C**) Bar graphs of the relative FPKM values of representative pro-inflammatory genes significantly downregulated upon TRIM24 overexpression, which was further validated by quantitative real-time PCR (**D**). (**E**) Chromatin immunoprecipitation sequencing (ChIP-Seq) of TRIM24-bound regions in NRVCMs with motif enrichment analysis (DREME/Tomtom), revealing enrichment of nuclear receptor motifs (RARα) and immune-related motifs recognized by STAT transcription factors. (**F**) Enrichment analysis of STAT-bound gene sets demonstrating preferential TRIM24 occupancy at STAT target loci. Statistical significance was determined using two-tailed Student’s *t* test. *Error bars* show means ± SEM. *, *p* < 0.05; **, p < 0.01; ***, p < 0.001.

Although TRIM24 does not directly bind DNA, its conserved functional domains allow it to act as a transcriptional coregulator, capable of modulating nuclear receptor-driven gene expression in both activating and repressive contexts^23^. To explore whether the transcriptomic changes observed upon TRIM24 overexpression are mediated through interactions with specific transcription factors, we conducted chromatin immunoprecipitation followed by sequencing (ChIP-Seq) in cardiomyocytes. Motif discovery using Discriminative Regular Expression Motif Elicitation (DREME) identified several short DNA motifs significantly enriched in TRIM24-bound regions. Subsequent comparison with known transcription factor binding profiles using the Tomtom algorithm revealed both expected and novel candidates, including the established nuclear receptor partner RARα and, notably, immune-related motifs such as those recognized by STAT transcription factors (Figure 2D). Additionally, genes known to be bound by STAT transcription factors showed high enrichment of TRIM24 (Figure 2E). These findings suggest that TRIM24 may exert part of its repressive effect on immune genes via cooperation with STAT-dependent regulatory elements.

### TRIM24 modulates STAT1/3 signaling in cardiomyocytes under inflammatory stimuli

Due to the observed strong effect of TRIM24 on inflammatory pathways, we next employed two complementary systems, LPS and IFNγ stimulation, to assess whether TRIM24 influences STAT1/3 activity across distinct inflammatory pathways. In unstimulated cells, TRIM24 overexpression reduced *Stat1a* and *Stat1b* transcript levels, while *Stat3* expression remained unchanged (Figure 3A-C). In contrast, TRIM24 knockdown significantly elevated expression of all three genes (Figure 3D-F), suggesting a baseline repressive effect of TRIM24 on STAT signaling. Following IFNγ stimulation, *Stat1a, Stat1b*, and *Stat3* transcripts were strongly induced. This induction was markedly blunted in TRIM24-overexpressing cells but further amplified when TRIM24 was silenced (Figure 3A-F). At the protein level, TRIM24 overexpression consistently reduced both basal and IFNγ-stimulated levels of total and phosphorylated STAT1/3 (Figure 3G-K), whereas TRIM24 knockdown enhanced their abundance and phosphorylation, particularly under IFNγ stimulation (Figure 3L-P). These results demonstrate that TRIM24 suppresses both gene expression and downstream activation of STAT1/3 in response to IFNγ.

**Figure 3.**
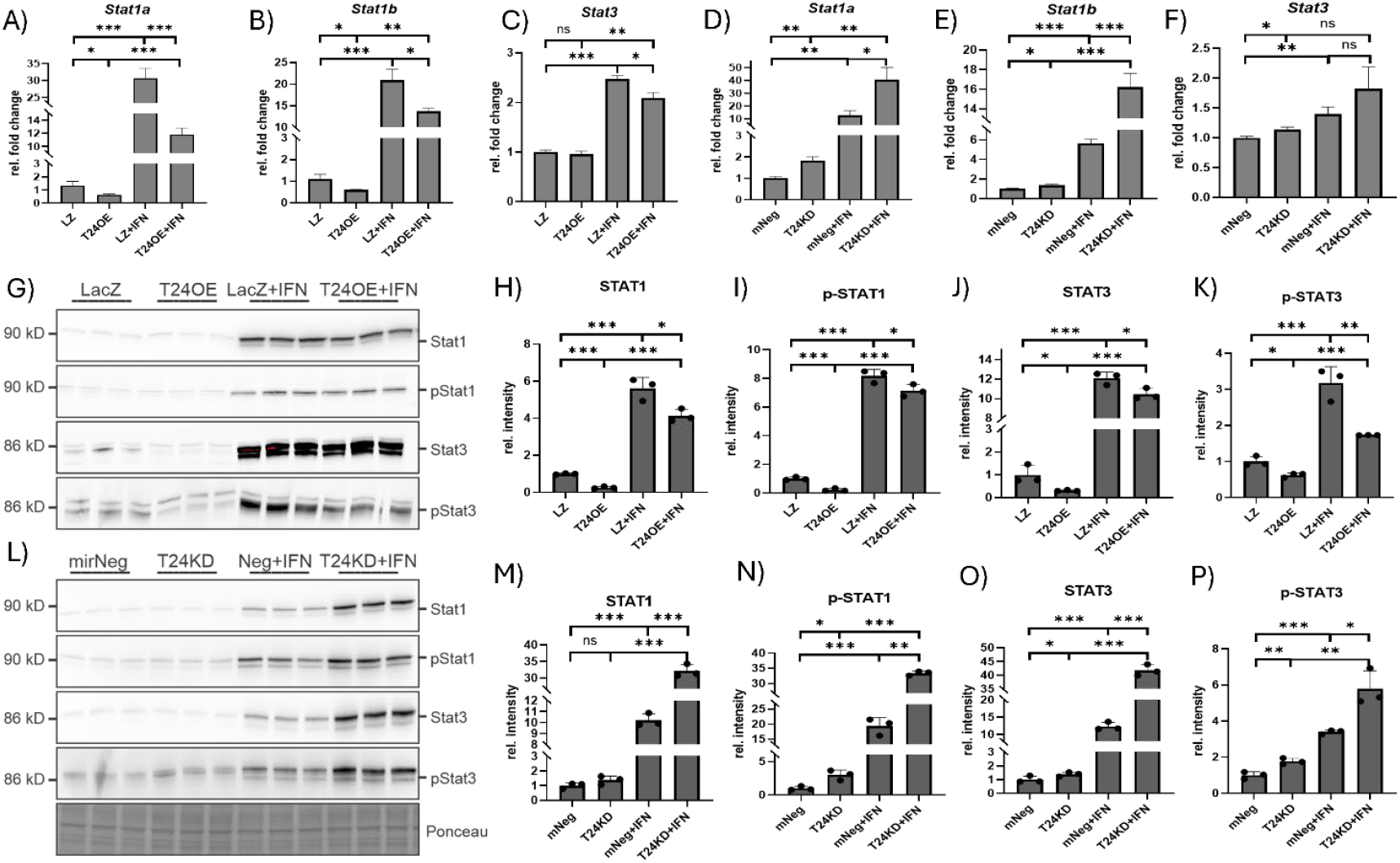
TRIM24 regulates STAT1/3 signaling in cardiomyocytes in response to IFNγ stimulation. Quantitative PCR analysis shows that TRIM24 overexpression reduces *Stat1a* (**A**), *Stat1b* (**B**), and *Stat3* (**C**) transcript levels in neonatal rat ventricular cardiomyocytes (NRVCMs) under both basal conditions and following interferon γ (IFNγ) stimulation. Conversely, TRIM24 knockdown upregulates *Stat1a* (**D**), *Stat1b* (**E**), and *Stat3* (**F**) expression at baseline, with further induction upon IFNγ treatment. Immunoblotting (**G**) and densitometric quantification (**H-K**) demonstrate that TRIM24 overexpression decreases total and phosphorylated STAT1 and STAT3 protein levels under resting and IFN-stimulated conditions. In contrast, immunoblotting (**L**) and corresponding densitometry (**M-P**) reveal that TRIM24 knockdown increases both total and phosphorylated STAT1 and STAT3, particularly following IFNγ stimulation. Statistical significance was determined using two-tailed Student’s *t* test. Error bars show means ± SEM. ns, non-significant; *, *p* < 0.05; **, *p* < 0.01; ***, *p* < 0.001.

LPS stimulation also increased STAT gene expression, though less robustly than IFNγ. TRIM24 overexpression significantly attenuated *Stat1a, Stat1b*, and *Stat3* transcript levels (Supplementary Figure 3A-C), while knockdown produced a trend toward higher expression, though statistical significance was not always reached (Supplementary Figure 3D-F). At the protein level, TRIM24 overexpression reduced STAT1, p-STAT1, and total STAT3 (Supplementary Figure 3G-K). Notably, unlike the IFNγ response, phosphorylated STAT3 was elevated in TRIM24-overexpressing cells upon LPS stimulation (Supplementary Figure 3G, 3K), indicating a context-dependent regulatory effect. TRIM24 knockdown increased STAT1 and p-STAT1 similar to the IFNγ condition, and also led to significant upregulation of p-STAT3, while total STAT3 remained largely unaffected (Supplementary Figure 3L-P).

Collectively, these data show that TRIM24 dampens STAT1/3 signaling in cardiomyocytes under both IFNΓ- and LPS-induced inflammation. While its repressive effect on STAT1 is consistent across conditions, regulation of STAT3 phosphorylation appears stimulus dependent.

### TRIM24 suppresses cardiomyocyte-driven macrophage migration

Building on the observation that TRIM24 modulates STAT1/3 activation and interferon-stimulated gene expression in cardiomyocytes, we next assessed the functional consequences of these molecular changes on immune cell behavior. Because cardiac inflammation involves paracrine signaling that guides immune cell recruitment, macrophage migration was used as a biological readout of cardiomyocyte-derived immune cues. To this end, transwell migration assay was performed using conditioned media from cardiomyocytes with TRIM24 overexpression, knockdown, or corresponding controls. Macrophages seeded in the upper chamber were allowed to migrate through porous membranes toward the respective supernatants. Quantitative analysis revealed that conditioned media from TRIM24-overexpressing cardiomyocytes significantly reduced macrophage recruitment, consistent with dampened cytokine signaling and repression of chemotactic interferon-stimulated genes. In contrast, supernatants from TRIM24-silenced cardiomyocytes promoted robust macrophage migration, reflecting an enhanced pro-inflammatory and chemotactic secretory profile (Figure 4A, 5B). These findings provide functional evidence that TRIM24 shapes the paracrine inflammatory capacity of cardiomyocytes by regulating immune cell recruitment.

**Figure 4:**
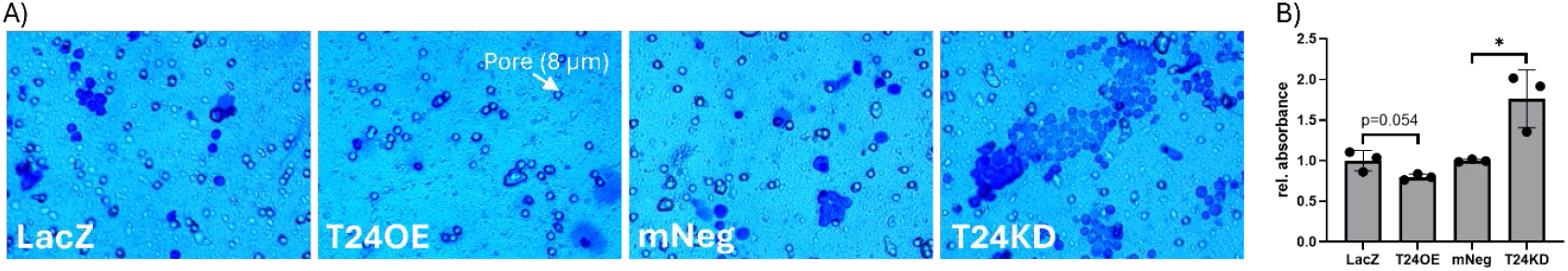
TRIM24 suppresses cardiomyocyte-driven macrophage migration. (**A**) Representative images from transwell migration assays in which macrophages migrated toward conditioned media derived from neonatal rat ventricular cardiomyocytes (NRVCMs) with TRIM24 overexpression, knockdown, or controls. (**B**) Quantification of migrated cells, presented as bar graphs, showing that conditioned media from TRIM24-overexpressing NRVCMs reduced macrophage migration, whereas supernatants from TRIM24 knockdown cells significantly enhanced migration. Data are presented as mean ± SEM. Statistical significance was determined using two-tailed Student’s *t*-test. *, *p* < 0.05.

**Figure 5:**
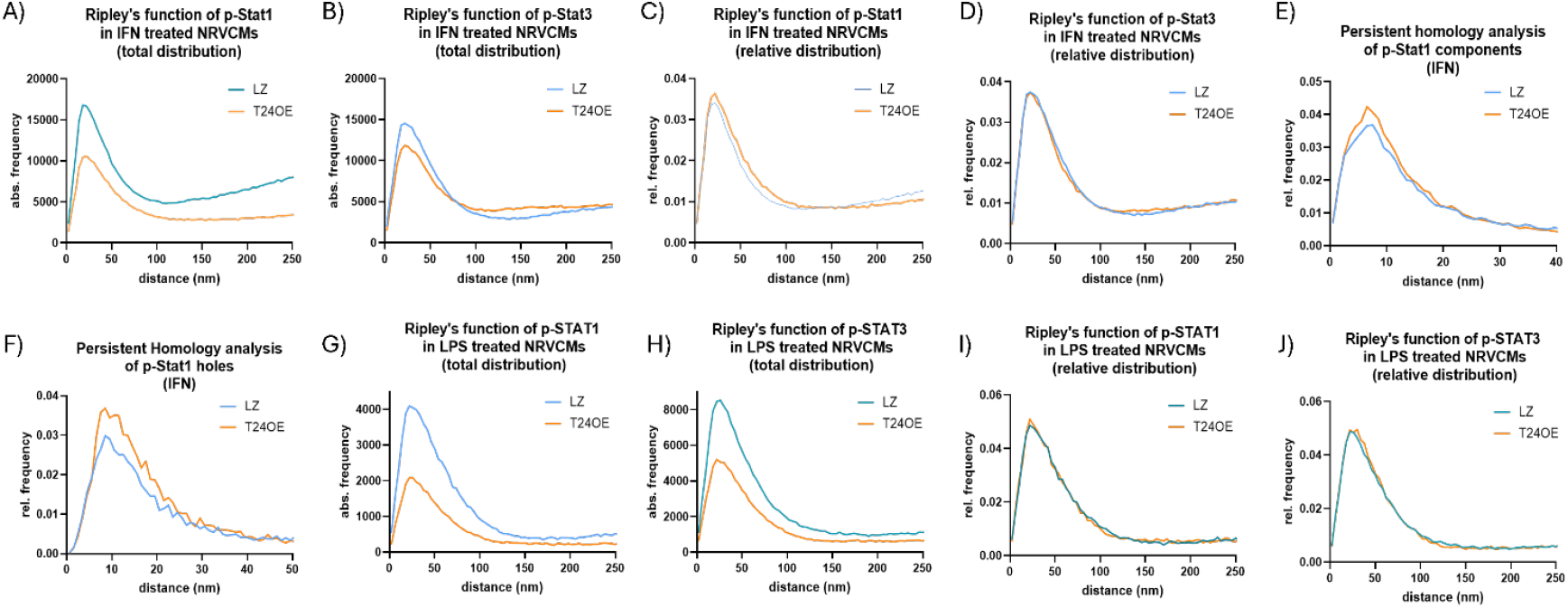
TRIM24 modulates nuclear abundance and nanoscale clustering of phosphorylated STAT1 and STAT3 under IFNγ and LPS stimulation. Super-resolution single-molecule localization microscopy was used to quantify nuclear organization of phosphorylated STAT1 (p-STAT1) and phosphorylated STAT3 (p-STAT3) in neonatal rat ventricular cardiomyocytes (NRVCMs) following interferon (IFNγ) or LPS stimulation. Spectral graphs depict (**A, B, G, H**) the absolute number of pair-wise distances of all blinking-event counts vs. distances between two points (absolute Ripley curves), showing reduced nuclear abundance of signals and clustering density (peak height and shape) of p-STAT1 and p-STAT3 in TRIM24-overexpressing cells under both stimuli. Relative Ripley analyses (**C, D, I, J**) normalized for total signal numbers reveal no or minimal reorganization of p-STAT3 and only modest alterations in p-STAT1 under IFNγ. Persistent homology analyses (relative frequency of components / holes vs. component / hole size represented by the distances of the respective bar end points) for p-STAT1 show minor changes in components (**E**) but increased maximum and width of hole distributions (**F**) under IFNγ, consistent with larger inter-cluster voids. Under LPS, notably, nuclear p-STAT3 levels were significantly reduced in TRIM24-overexpressing cells after LPS stimulation (**G**).

### TRIM24 modulates nuclear clustering and nanoscale topology of phosphorylated STAT1 and STAT3 under inflammatory stress

We observed that TRIM24 regulates both the expression and phosphorylation of STAT1 and STAT3 in cardiomyocytes (see Figures 2 and 3). Moreover, ChIP-seq motif enrichment analysis revealed a significant overrepresentation of STAT consensus motifs within TRIM24-bound genomic regions (see Figures 1D, 1E), implicating TRIM24 in the modulation of STAT-dependent transcriptional programs. Given that transcriptional output is strongly influenced by the spatial clustering and higher-order nuclear topology of transcription factors, to examine whether TRIM24 alters the subnuclear distribution of activated STAT proteins, we performed super-resolution single-molecule localization microscopy of phosphorylated forms of STAT1 (p-STAT1) and STAT3 (p-STAT3) in NRVCMs after IFNγ or LPS stimulation. Spatial patterns were quantified using total blinking-event counts, absolute Ripley functions, relative Ripley functions (normalized for event count), and persistent homology (component and hole analyses). Representative images and quantitative analyses are shown in Figure 5 and Supplementary Figure 4.

Under IFNγ stimulation, TRIM24 overexpression produced a clear reduction in nuclear p-STAT1 and p-STAT3 signal, as evidenced by lower total blinking-event counts and reduced absolute Ripley values (Figures 5A, 5B). The absolute Ripley curves for TRIM24-overexpressing cells approached the random-distribution bisector more slowly than controls, consistent with reduced local density of phosphorylated STAT molecules (Figures 5A, 5B). When expression differences were accounted for by relative Ripley analysis, p-STAT3 spatial organization was not significantly altered, while p-STAT1 showed only a modest, but measurable, deviation from control (Figures 5C, 5D). Persistent homology component analysis of p-STAT1 revealed only a minor difference between control and TRIM24OE conditions (Figures 5E), whereas the holes analysis for p-STAT1 showed an increased maximum and a broader width in TRIM24-overexpressing cells (Figures 5F), indicating larger inter-cluster gaps. Persistent homology metrics for p-STAT3 under IFNγ showed no significant differences (Supplementary Figures 4A, 4B).

Under LPS stimulation, TRIM24 overexpression likewise reduced total nuclear blinking events which resulted in reduced absolute Ripley values for both p-STAT1 and p-STAT3 (Figures 5G, 5H). However, after normalization of the pairwise distance frequency distributions (relative Ripley curves) and in topological (persistent homology) analyses, neither phospho-STAT species displayed significant reorganization between control and TRIM24OE (Figures 5I, 5J; Supplementary Figures 4C-F). Notably, microscopy detected a significant reduction of the nuclear p-STAT3 pool in TRIM24-overexpressing cells following LPS (Figures 5G), a change that was not apparent in bulk biochemical assays (see Figure 2), indicating a selective effect on the nuclear-localized activated STAT3 population.

Direct comparison of stimuli showed that IFNγ treatment produced substantially more blinking events for both p-STAT1 and p-STAT3 than LPS (Figures 5A vs 5G; Figures 5B vs 5H). Under IFNγ, the Ripley function for p-STAT1 transitioned towards random distribution at shorter distances, whereas in the LPS condition p-STAT1 remained more dispersed with very low clustering frequency beyond ~150 nm (Figures 5A, 5G). A similar but less pronounced stimulus-dependent pattern was observed for p-STAT3 (Figures 5B, 5H). Holes analysis of p-STAT1 further showed that IFNγ-treated samples reached a maximal hole signal earlier but at a lower amplitude than LPS samples (Figures 5F). Persistent homology analyses for p-STAT3 did not reveal comparable stimulus-dependent topology changes. In summary, TRIM24 overexpression reduces nuclear abundance of activated STAT1 and STAT3 under both IFNγ and LPS stimulation and selectively alters the nanoscale topology of p-STAT1 (larger inter-cluster v`oids) under IFNγ. These imaging results, which specifically report on nuclear, clustered pools of activated STATs, complement the bulk biochemical measurements and suggest TRIM24 influences both the nuclear abundance and nanoscale organization of activated STAT complexes.

### TRIM24 suppresses STAT1/3 via its chromatin reader domain rather than proteasomal degradation

Transcriptomic analyses indicated that TRIM24 reduces STAT1 and STAT3 expression at the transcriptional level. However, because TRIM24 also harbors E3 ubiquitin ligase activity, it remained unclear whether the decrease in protein abundance might additionally involve post-translational degradation. To test this possibility, cardiomyocytes were treated with MG132, a proteasome inhibitor, that blocks ubiquitin-mediated protein turnover. Proteasome inhibition did not restore STAT1 or STAT3 levels in TRIM24-overexpressing cells, indicating that TRIM24 does not promote their degradation via ubiquitination but instead acts upstream, most likely at the level of transcriptional regulation (Figure 6 A, 6B & Supplementary Figure 5A, 5B). We next asked whether chromatin binding through the PHD-bromodomain module of TRIM24 is required for STAT repression. TRIM24-overexpressing cardiomyocytes were treated with IACS-9571, a selective inhibitor of the TRIM24 bromodomain. Pharmacological inhibition partially restored STAT1 and STAT3 expressions, strongly suggesting that the chromatin reader function of TRIM24 is critical for their transcriptional repression (Figure 6 A, 6C & Supplementary Figure 5A, 5C)

To investigate how bromodomain inhibition alters spatial organization of TRIM24, we performed single-molecule localization microscopy. Quantitative analyses revealed decreased nuclear clustering of TRIM24 and a concomitant increase in cytosolic organization upon inhibitor treatment. While nuclear topological changes were minimal, cytosolic clusters became more prominent, indicating a redistribution of TRIM24 from the nucleus to the cytoplasm (Figure 6D-G). Together, these findings identify TRIM24 as a transcriptional repressor of STAT1 and STAT3 in cardiomyocytes. This function depends on its PHD-bromodomain chromatin reader module rather than its E3 ubiquitin ligase activity. Inhibition of the bromodomain not only relieves STAT repression but also reshapes subcellular organization of TRIM24, highlighting the central role of chromatin interactions in its regulatory activity under inflammatory stress.

**Figure 6:**
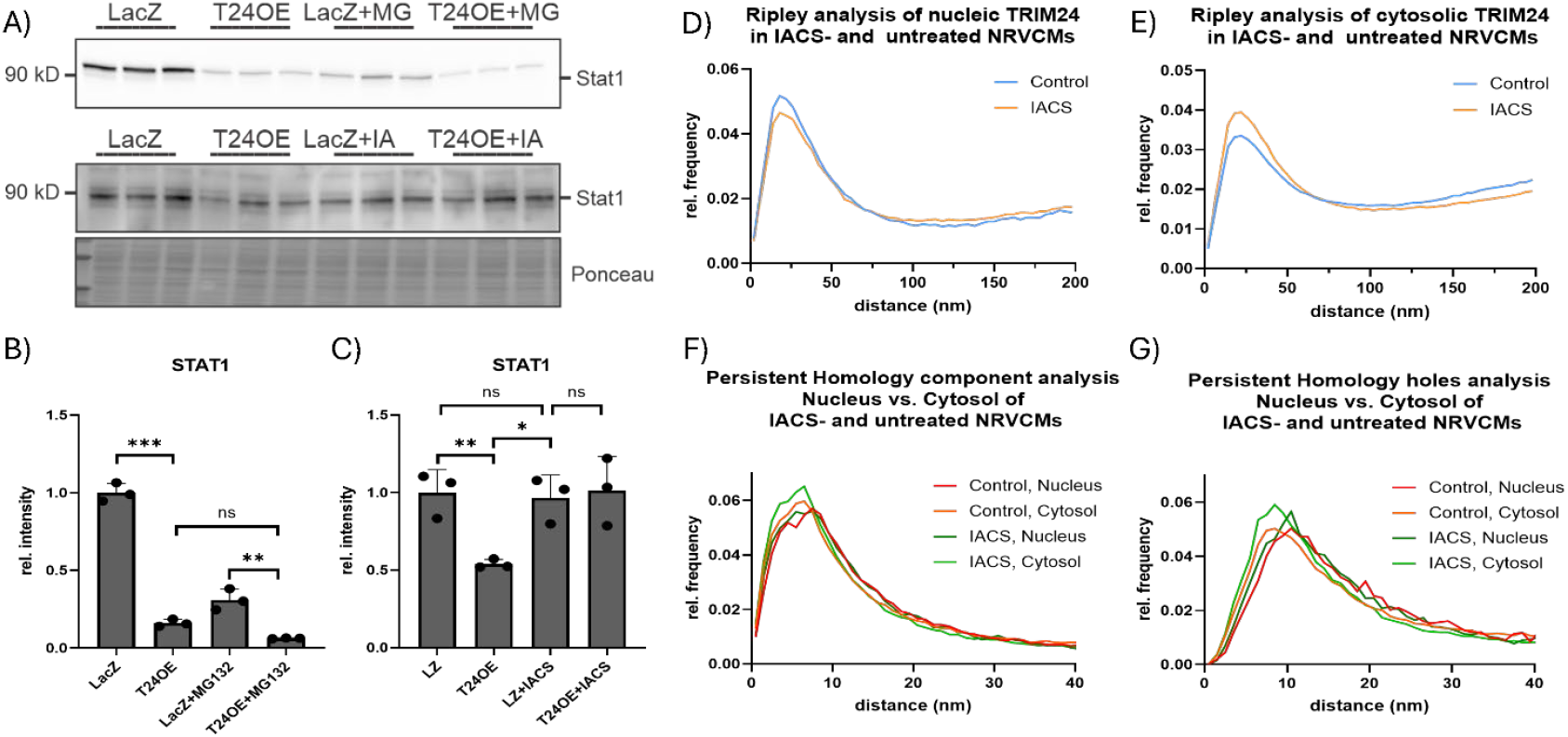
TRIM24 represses STAT1 and STAT3 via its chromatin reader domain rather than proteasomal degradation. Immunoblotting (**A**) and densitometric analysis show that proteasomal inhibition by MG132 does not prevent TRIM24-mediated downregulation of STAT1 (**B**), whereas treatment with IACS attenuates the TRIM24-induced decrease in STAT1 protein levels (**C**). Super-resolution single-molecule localization microscopy was used to quantify nuclear and cytosolic organization of TRIM24 in neonatal rat ventricular cardiomyocytes (NRVCMs) following IACS-5971 treatment. Spectral graphs depict the absolute number of pair-wise distances of all blinking-event counts vs. distances between two points (absolute Ripley curves), showing reduced nuclear (**D**) while increased cytosolic (**E**) abundance and clustering density of TRIM24 in IACS-5971 treated cells. Persistent homology analyses show minor changes in components (**F**) and width of hole distributions (**G**) in the nucleus compared to the cytosol.

## Discussion

The present study identifies a role for TRIM24 in transcriptional regulation of immune signaling in cardiomyocytes, exerting a predominantly repressive effect on interferon-stimulated genes and the STAT1/3 axis. Integrative analyses combining RNA-seq, proteomics, and ChIP-seq consistently pointed to a dampening of cytokine- and interferon-related pathways upon TRIM24 overexpression. These molecular changes were functionally translated into a decreased capacity of TRIM24 overexpressing cardiomyocytes to recruit macrophages via paracrine signaling, emphasizing the physiological relevance of TRIM24-dependent immune suppression in the cardiac environment. Given the already mentioned cell type-specific duality of TRIM24 function, this study is the first to demonstrate that TRIM24 exerts a suppressive effect on immune signaling in cardiomyocytes. Our findings further extend these observations by establishing that TRIM24 is a predominantly nuclear protein in cardiomyocytes, enriched within the chromatin fraction and markedly upregulated in healthy myocardium compared to failing or ischemic hearts. This subcellular localization pattern, corroborated in cardiomyocytes isolated from both sham and LAD-ligated mice, highlights TRIM24 as a chromatin-associated safeguard whose nuclear depletion accompanies cardiac stress and inflammation. Together with its repressive effects on interferon signaling, these data position TRIM24 as a key nuclear modulator maintaining immune quiescence in the myocardium.

The inhibitory regulation of STAT1 and STAT3 by TRIM24 was confirmed at the protein level in NRVCMs following both IFNγ and LPS stimulation. Immunoblot analysis consistently demonstrated TRIM24-dependent regulation of total STAT1 and STAT3 levels. Overexpression of TRIM24 resulted in their downregulation, whereas knockdown led to increased levels, independent of whether the cells were stimulated with IFNγ or LPS. This suppressive effect was also observed for the phosphorylated forms of STATs. An unexpected finding was the significant increase in p-STAT3 levels upon LPS stimulation in TRIM24OE cells, which appears to contradict the overall trend. A possible explanation for this could be TRIM24-mediated modulation of upstream JAK or tyrosine kinase activity, thereby influencing phosphorylation dynamics independently of total STAT protein levels. Supporting this notion, previous studies have shown that TRIM24 depletion reduces JAK1 and JAK2 phosphorylation in macrophages following LPS stimulation, indicating that TRIM24 may regulate the JAK/STAT pathway by modulating JAK2 activity upstream^38^. However, direct interactions between TRIM24 and JAKs have yet to be demonstrated. Alternatively, the involvement of NF-κB, a well-known regulator of STAT3 phosphorylation^39^ and itself modulated by TRIM24^40^, may play a role under these specific proinflammatory conditions. TRIM24 has been shown to influence NF-κB target gene expression, thereby promoting IL-6 transcription through chromatin interactions or co-activator functions. Elevated IL-6 production could, in turn, enhance STAT3 phosphorylation via the gp130/JAK pathway ^41^. Such IL-6/STAT3 positive feedback loops have been described in tumor models and may also be reinforced by TRIM24 in the cardiac context. Additionally, TRIM24 overexpression might suppress IL-10 production or delay its induction in LPS-stimulated cardiomyocytes^16^. Since IL-10 normally limits IL-6-induced STAT3 activation via SOCS3, reduced IL-10 levels could prolong STAT3 phosphorylation^42^. Furthermore, by repressing the IFN/STAT1 axis, TRIM24 may lower SOCS1 and SOCS3 expression, further weakening the negative feedback regulation of the JAK/STAT3 pathway. Collectively, this imbalance could explain the observed increase in p-STAT3 levels under LPS stimulation, despite TRIM24’s generally repressive role.

ChIP-seq analysis revealed distinct TRIM24 binding enrichments at several immune-regulatory genes highlighting the variable binding patterns of TRIM24, ranging from sharp, well-defined peaks, such as at the *Stat6* locus, to broader enrichments observed at *Irf7* and *Ifit* genes. This diversity points towards TRIM24’s versatile regulatory roles, particularly in immune signaling. Motif analysis confirmed the presence of the known RARα motif, supporting a previously proposed mechanism of STAT regulation described by Tisserand et al^17^. Interestingly, novel and previously unreported STAT1/STAT3 motifs were identified. Given that STAT1 and STAT3 share similar binding motifs and considering that TRIM24 has been shown to bind and stabilize STAT3 chromatin interactions^18^, this may indicate a potential regulatory interaction. Additionally, the transcription factor motif NHLH2 was detected. The NHLH2:STAT3 heterodimer is a known transcriptional regulator, with NHLH2 binding to E-box motifs located adjacent to STAT3 binding sites^43^. This suggests a possible functional interplay between TRIM24, STAT3, and NHLH2 in controlling immune-related gene expression. Moreover, the clear enrichments observed at *Ifit* and *Irf* genes imply that TRIM24 may also modulate STAT1-driven transcriptional programs, although further experimental validation will be required to confirm this.

The observation that inhibition of the TRIM24 bromodomain by IACS-9571 alleviates TRIM24-mediated repression of STAT1 and STAT3 aligns with previous findings showing that TRIM24 recruits the transcription factor RARα to the ORM2 promoter through its interaction with H3K27ac^44^. This suggests that a similar mechanism may also be operative in cardiomyocytes. However, in the context of ORM2, TRIM24 functions as a co-activator, rather than acting as a repressor of RARα, as it does here. Since the same primary transcription factor is involved in both contexts, this raises a fundamental question: how can TRIM24 switch between functioning as a transcriptional activator and a repressor? Neither the presence of a specific histone modification nor the identity of the main transcription factor alone seems sufficient to explain this duality. Instead, it points to a more complex regulatory mechanism, likely influenced by additional cofactors, the surrounding chromatin landscape, or even the spatial organization within the nucleus. One possible explanation is that IACS-9571 not only blocks the bromodomain of TRIM24 but also interferes with its SUMOylation, a post-translational modification known to affect protein interactions and functional activity. Such context-dependent regulatory plasticity underscores TRIM24’s ability to fine-tune transcriptional output in a stress- and chromatin-dependent manner, rather than functioning as a binary on/off switch for gene expression. Taken together, the ChIP-seq and Western blot data presented in this study demonstrate for the first time that TRIM24 plays a key role as suppressor of immune signaling in cardiomyocytes.

Furthermore, the detection of STAT motifs in the ChIP-seq dataset, along with TRIM24 binding enrichments at ISG loci, further suggests that TRIM24 may have a more complex regulatory role in immune signaling than previously understood. Single-molecule localization microscopy added a spatial dimension to these findings. TRIM24 overexpression reduced nuclear pools of p-STAT1 and p-STAT3 under both IFNγ and LPS stimulation, and selectively altered the nanoscale topology of p-STAT1 under IFNγ. Notably, microscopy revealed decreased nuclear p-STAT3 after LPS stimulation, a change not apparent by bulk western blotting, suggesting that TRIM24 may influence STAT3 nuclear import or compartmentalization. Together with its enrichment in the nuclear fraction of cardiomyocytes, these findings support a model in which TRIM24 not only represses STAT transcriptional programs but also modulates the higher-order nuclear organization of signaling complexes. Such nuclear architectural regulation, coupled with transcriptional repression, indicates that TRIM24 coordinates both the abundance and spatial organization of STAT complexes.

Beyond mechanistic insight, these findings carry important pathophysiological implications. The loss of nuclear TRIM24 observed in failing and ischemic hearts suggests that its depletion may relieve repression of interferon-responsive genes, fueling chronic inflammatory activation. This aligns with transcriptomic data from heart failure patients showing persistent activation of interferon-stimulated genes and STAT-driven networks. Thus, maintaining or restoring TRIM24 function could represent a novel strategy to restrain sterile inflammation in cardiac disease. Together, these findings position TRIM24 as a dynamic regulator of immune signaling in cardiomyocytes, integrating transcriptional, post-transcriptional, and spatial control. By repressing STAT-driven inflammatory programs while modulating nuclear STAT architecture, TRIM24 shapes the paracrine output of cardiomyocytes and thereby regulates macrophage recruitment. Given its dual roles as a chromatin reader and transcriptional coregulator, TRIM24 emerges as a potential therapeutic target to modulate cardiac inflammation. Targeting TRIM24 or its bromodomain could provide new strategies to fine-tune detrimental immune responses in cardiovascular disease while preserving essential homeostatic functions.

### Significance and clinical importance

Our study uncovers TRIM24 as a previously unrecognized transcriptional repressor of immune signaling in cardiomyocytes. By suppressing interferon-stimulated genes and modulating STAT1/3 pathways at both molecular and spatial levels, TRIM24 shapes the inflammatory output of cardiomyocytes and limits macrophage recruitment. These findings extend the functional repertoire of TRIM24 beyond cancer biology into cardiovascular immunology, highlighting it as a potential therapeutic target for fine-tuning detrimental cardiac inflammation. Pharmacological inhibition of the TRIM24 bromodomain reversed STAT repression, underscoring its druggable potential and paving the way for therapeutic interventions aimed at mitigating immune-driven cardiac injury, such as viral myocarditis or cancer therapy–induced cardiotoxicity.

### Limitations of the Study

While this work provides multi-level evidence that TRIM24 suppresses cardiomyocyte immune signaling, several limitations should be acknowledged. First, the study is confined to in vitro neonatal rat ventricular cardiomyocytes, and in vivo validation in models of cardiac inflammation is required to confirm physiological relevance. Second, although our data implicate chromatin binding via the PHD-bromodomain in TRIM24-mediated repression, the precise cofactors and chromatin landscapes that determine its activator-versus-repressor duality remain unresolved. Third, the discrepancy between bulk biochemical assays and super-resolution imaging for STAT3 highlights the complexity of TRIM24 regulation, but the exact mechanisms governing STAT3 nuclear import and compartmentalization need further investigation. Finally, our study does not directly address the potential systemic consequences of TRIM24 modulation, particularly in immune or non-cardiac cells, which could impact the translational feasibility of TRIM24-targeted therapies.

## Declarations

### Ethics approval and consent to participate

Not applicable

### Consent for publication

All authors declare consent for publication

### Availability of data and materials

All data underpinning the findings of this study are comprehensively provided within the manuscript or in supplementary data file. The mass spectrometry proteomics data have been deposited to the ProteomeXchange Consortium via the PRIDE partner repository with the dataset identifier PXD062686.

### Authors’ contributions

M.N., M.H., M.K., and A.Y.R. conceived and designed the experiments; M.N., A.B., A.D., L.N., T.B., F.S., E.H., N. S., M. K., and A.Y.R. performed the experiments and/or analyzed the data; E.H., M.H., M.K., N.F., and A.Y.R. contributed reagents/materials/analysis tools; M.N., and A.Y.R. wrote the manuscript; M.N., A.B., A.D., E.H., M.K., N.F., and A.Y.R. revised the manuscript. All authors have read and agreed to the submitted manuscript.

## Acknowledgements

We thank Wojciech Wilk and Luisa Lange for excellent technical support as well as Vishnu Mukund Dhople for mass spectrometric data acquisition.

