## supplement table and supplementary figures for "TRIM24 preserves cardiomyocyte immune quiescence by repressing interferon/STAT signaling"

**Supplementary Table 1: Oligonucleotide sequences of primers used for the quantitative real-time PCR. hs – Homo sapiens; rn – Rattus norvegicus**

| Primer name | Sequence | Gene |
|---|---|---|
| hs_TRIM24_ rt-fw | 5'-AGCCTAGCTCAATTACGGCTC-3' | TRIM24 |
| hs_TRIM24_ rt-rev | 5'-GCGGTTGCTGATGAGAGATGG-3' | |
| rn_TRIM24_ rt-fw | 5'-CAACAGGCCATAAAACAGTGGC-3' | TRIM24 |
| rn_TRIM24_ rt-rev | 5'-GCACTTGTGATCGTGGGACT-3' | |
| rn_ Ifit2_ rt-fw | 5'-TCATGAGTACAGCCAGTAAGGAA-3' | IFIT2 |
| rn_ Ifit2_ rt-rev | 5'-ACTCGTCTTCTGCTCTCAGG-3' | |
| rn_ Stat1a_ rt_ fw | 5'-AAAGGAAGCACCAGAACCGAT -3' | STAT1a |
| rn_ Stat1a_ rt_ rev | 5'-CTCTGGAGACATGGGAAGCA -3' | |
| rn_ Stat1b_ rt_ fw | 5'-AAAGGAAGCACCAGAACCGAT-3' | STAT1b |
| rn_ Stat1b_ rt_ rev | 5'-AGGTTCTCAACAAGCCAGTCT-3' | |
| rn_ Stat3_ rt_ fw | 5'-GACATTCCCAAGGAGGAGGC-3' | STAT3 |
| rn_ Stat3_ rt_ rev | 5'-GGCAGCACTACCTGGGTC-3' | |
| rn_ RPL32_ rt_ fw | 5'-GGTGGCTGCCTTTACG-3' | RPL32 |
| rn_ RPL32_ rt_ rev | 5'-CCGCACCCTGCAATGC-3' | |

### Supplementary Figures

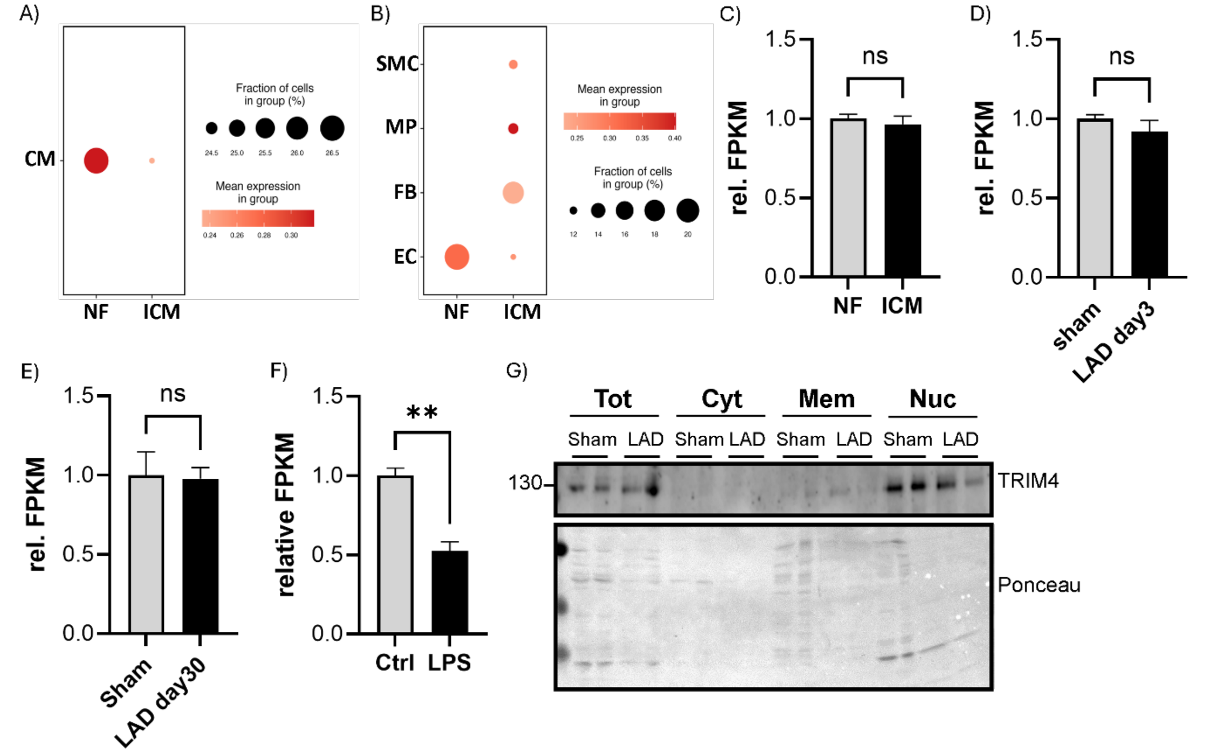


**Supplementary Figure 1. TRIM24 is a cardiomyocyte-enriched nuclear factor reduced in human ischemic cardiomyopathy and regionally regulated after myocardial infarction.** Dot plots of TRIM24 transcript levels in cardiomyocytes (**A**) and other cardiac cell types (**B**) from single-nucleus RNA sequencing dataset GSE121893, showing reduced expression in ischemic cardiomyopathy (ICM) compared with non-failing (NF) hearts. (**C**) Bulk RNA-seq analysis of TRIM24 levels in left ventricular tissue from non-failing (NF) and ischemic cardiomyopathy (ICM) patients using dataset GSE141910, showing no significant change. Bulk RNA-seq data of Trim24 expression in mouse hearts following acute (3 days; **D**) or chronic (30 days; **E**) LAD ligation showing no significant difference compared with sham controls. (**F**) Trim24 mRNA levels significantly reduced in mouse hearts 24 h after lipopolysaccharide (LPS) treatment (dataset GSE185754). (**G**) Subcellular fractionation and immunoblotting of cardiomyocytes showing TRIM24 protein predominantly detected in nuclear and total lysate fractions, with minimal presence in membrane and cytoplasmic fractions. Statistical significance was determined using two-tailed Student's *t* test. Error bars represent mean ± SEM. ns, non-significant; **, $p < 0.01$.

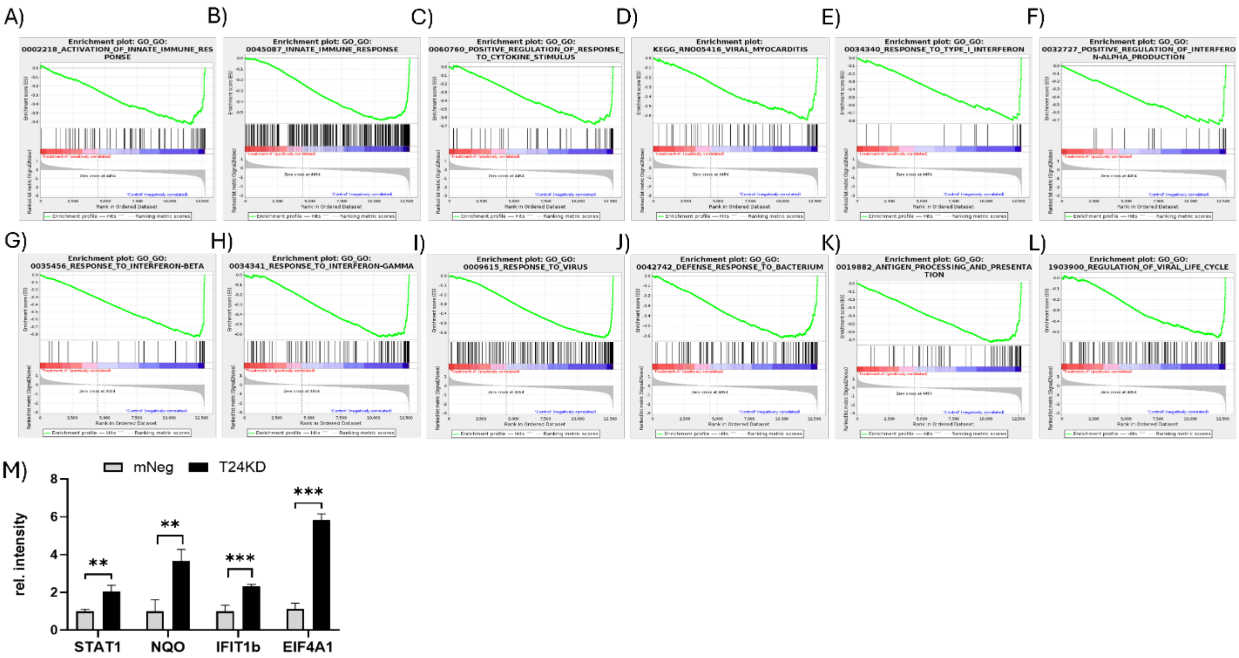


**Supplementary Figure 2: TRIM24 reprograms the cardiomyocyte immune transcriptome and targets STAT-related regulatory networks.** (**A-L**) GSEA-plots indicating the effects of TRIM24 overexpression on immune response related pathways**.** All of them exhibit a clear negative Enrichment Score. (**M**) Bar graph depicting upregulation of few of the proteins involved in Stat1 signaling in NRVCMs when TRIM24 is knocked-down, determined by mass-spectrometry.

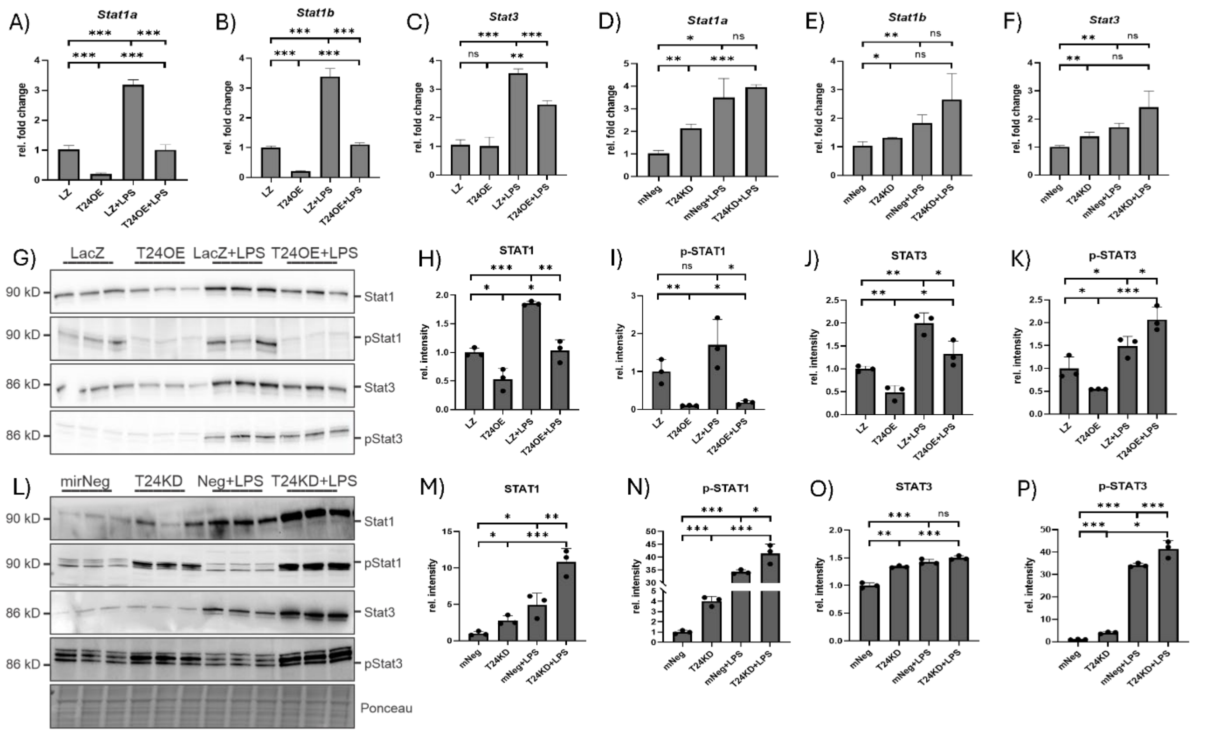


**Supplementary Figure 3. TRIM24 regulates STAT1/3 signaling in cardiomyocytes in response to LPS stimulation.** Quantitative PCR analysis shows that TRIM24 overexpression suppresses *Stat1a* (**A**), *Stat1b* (**B**), and *Stat3* (**C**) transcript levels in neonatal rat ventricular cardiomyocytes (NRVCMs) under both basal and LPS-stimulated conditions. TRIM24 knockdown produced a trend toward higher expression of *Stat1a* (**D**), *Stat1b* (**E**), and *Stat3* (**F**), although statistical significance was not consistently achieved. Immunoblotting (**G**) and densitometric quantification (**H-K**) demonstrate that TRIM24 overexpression reduces STAT1, phosphorylated STAT1, and total STAT3 protein levels following LPS treatment. Conversely, immunoblotting (**L**) and densitometry (**M-P**) show that TRIM24 knockdown elevates STAT1 and phosphorylated STAT1, while also significantly enhancing phosphorylated STAT3 levels, with total STAT3 remaining largely unchanged. Statistical significance was determined using two-tailed Student's *t* test. *Error bars* show means ± SEM. ns, non-significant; *, $p < 0.05$; **, $p < 0.01$; ***, $p < 0.001$.

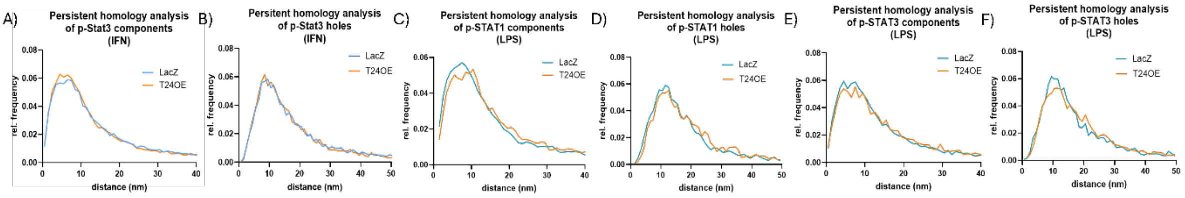


**Supplementary Figure 4: Extended persistent homology analyses of nuclear p-STAT1 and p-STAT3 under IFN and LPS stimulation.** Persistent homology metrics for p-STAT3 under IFN stimulation, showing no significant differences between control and TRIM24-overexpressing NRVCMs in either component counts (**A**) or hole distributions (**B**). Persistent homology analyses for p-STAT1 (**C, D**) and p-STAT3 (**E, F**) under LPS stimulation, demonstrating no significant changes in components (**C, E**) or holes (**D, F**) upon TRIM24 overexpression.

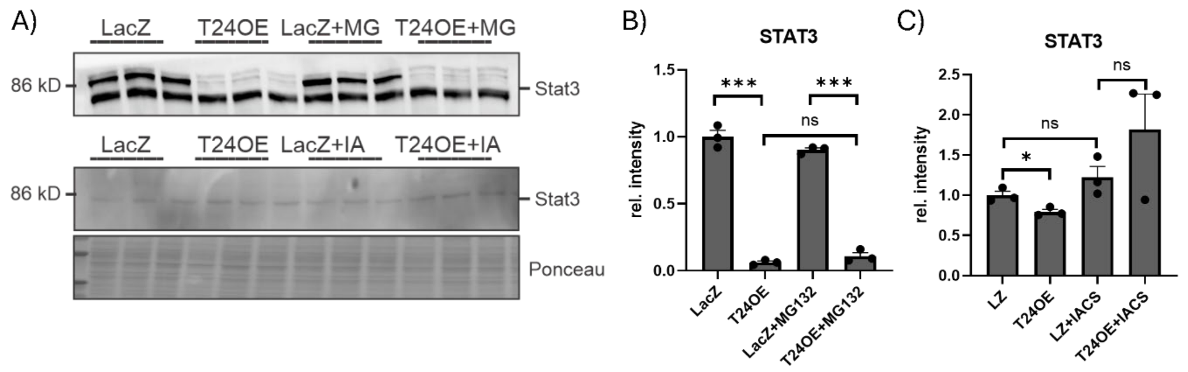


**Supplementary Figure 5: TRIM24 suppresses STAT1/3 via its chromatin reader domain rather than proteasomal degradation.** Immunoblotting (**A**) and densitometric analysis show that proteasomal inhibition by MG132 does not prevent TRIM24-mediated downregulation of STAT1 (**B**), whereas treatment with IACS attenuates the TRIM24-induced decrease in STAT1 protein levels (**C**).
